# Extracellular microbes are required for mosquito development even in the presence of *Wolbachia*

**DOI:** 10.1101/2024.12.03.626537

**Authors:** Javier Serrato-Salas, Danai Bemplidaki, Ivan Roger, Yanouk Epelboin, Mathilde Gendrin

## Abstract

*Wolbachia* is an endosymbiotic bacterium infecting a wide array of invertebrates that gained attention for its potential to curb the transmission of vector-borne diseases. Its capacity to colonize arthropod populations is generally driven by vertical transmission and reproductive manipulation. In some insect species, *Wolbachia* additionally became an essential nutritional symbiont, providing vitamins to its host. As mosquito larvae require microbe-derived vitamins for development, we studied whether such a support of *Wolbachia* would exist in mosquitoes but be masked by the presence of other microbes. We chose *Culex quinquefasciatus* species to address this question, as it is highly colonized with *Wolbachia*. We developed a method to produce *Culex quinquefasciatus* devoid of extracellular microbiota and demonstrated that *Wolbachia* alone is insufficient to support larval development. Using transient colonization with *Escherichia coli*, we managed to produce adult *Culex quinquefasciatus* harboring *Wolbachia* only. When curbing *Wolbachia* infection of these *E. coli*-colonized larvae via tetracycline treatment, we obtained a higher larval development. Together, our data indicate that *Wolbachia* does not support development but rather acts here as a metabolic burden, and that *E. coli* is sufficient for development success even in a species that grows in “dirty” water. This opens the way towards gnotobiology studies in *Culex quinquefasciatus* and highlights the complex relationships between *Wolbachia* and its mosquito host.

**Author summary:** *Wolbachia* is a bacterium infecting many invertebrates that gained attention for its potential to curb the transmission of vector-borne diseases, such as dengue. In some insect species, *Wolbachia* provides vitamins to its host. As mosquito larvae require microbe-derived vitamins for development, we studied whether such a support of *Wolbachia* would exist in mosquitoes but be masked by the presence of other microbes. We chose *Culex quinquefasciatus* species to address this question, as it is highly colonized with *Wolbachia*. We developed a method to produce *Culex quinquefasciatus* devoid of extracellular microbiota and demonstrated that *Wolbachia* alone is insufficient to support larval development. We transiently colonized these larvae with *Escherichia coli* to rescue larval development and to produce adults harboring *Wolbachia* only. When curbing *Wolbachia* infection of these *E. coli*-colonized larvae via tetracycline treatment, we obtained a higher larval development. Together, our data indicate that *Wolbachia* does not support development but rather acts here as a metabolic burden, and that *E. coli* is sufficient for development success even in a mosquito species that grows in “dirty” water.

## Introduction

Mosquito-borne diseases are a worldwide health threat. New vector control tools that are safe for people and the environment are needed. One efficient approach uses the release of *Aedes* mosquitoes infected with the endosymbiotic bacterium *Wolbachia* to suppress mosquito populations, ultimately reducing the incidence of mosquito-borne diseases (1). This approach has allowed significant progress in reducing dengue incidence in high burden settings worldwide (2,3).

*Wolbachia* is a genus of intracellular alphaproteobacteria that infect many invertebrate species. It has attracted particular interest due to its diverse effects on hosts. In arthropods, *Wolbachia* is present in approximately 65 % of terrestrial species (4); and half of aquatic species (5). While its interaction with nematodes is clearly mutualistic, contributing to worm nutrition, its role in arthropods is generally parasitic, as it manipulates host reproduction for its own benefit (6–8). Within arthropods, it is highly concentrated in female germlines, enabling vertical transmission while simultaneously acquiring resources for survival and growth (8).

Among the reproductive manipulation phenomena, cytoplasmic incompatibility (CI) is most common. In CI, infected males mating with uninfected females produce embryonic death, while they can fertilize infected females, resulting in viable offspring. This increases the relative competitiveness of infected females within the population, promoting *Wolbachia* transmission (9). When two mosquito populations harbor different *Wolbachia* strains, bidirectional CI can occur. *Wolbachia* also induces other phenotypic changes, often increasing the populatiońs proportion of females, which further enhances its spread. These changes include: a) induction of parthenogenesis, leading to female offspring; b) male killing, where *Wolbachia* infection in mated females selectively eliminates developing male embryos by manipulating the sex-determination system; and c) feminization, where males develop female characteristics due to hormone manipulation (10).

Recently, two hemipteran species have been found to maintain an unexpected nutritional link with *Wolbachia.* In bedbugs, *wCle* (*Wolbachia* of *Cimex lectularius*) promotes nymph development, lifespan and fecundity by producing riboflavin and biotin (8,11,12). In two planthoppers species, *Wolbachia*-cured insects are sterile, but their fecundity levels are rescued by experimental reinfection with *wLug* (*Wolbachia* of *Nilaparvata lugens)* and *wStri (Wolbachia* of *Laodelphax striatellus*), respectively (13).

*Wolbachia* has been found naturally in some dipterans, such as mosquitoes and flies (14–16). This includes the common domestic mosquitoes *Culex quinquefasciatus* and *Culex pipiens* where *Wolbachia* is highly prevalent (17), in contrast to other mosquito species, notably *Aedes aegypti* and *Anopheles* species, where *Wolbachia* has rarely been detected (18–20).

*Culex* mosquitoes are vectors of multiple medically significant arboviruses, such as West Nile virus, Japanese encephalitis virus, Usutu virus, Saint Louis encephalitis virus, Western and Eastern equine encephalitis viruses (21–24). In addition to viruses, *Cx* mosquitoes can also transmit nematodes responsible for lymphatic filariasis and protists that cause avian malaria (25,26).

Another known role for *Wolbachia* in dipterans is increased resistance to viral replication in flies (27) as well as in transinfected mosquitoes (28,29). In *Culex* mosquitoes, several studies have shown that, except for bidirectional cytoplasmic incompatibility, the impact of *Wolbachia* on life history traits is low or moderate. *Wolbachia-*cured mosquitoes exhibit an increased and a delayed egg production, a minor decrease in lifespan, and a slight decrease in resistance to mosquitocidal bacterial strains (30,31). *Wolbachia* in *Culex* mosquitoes exhibits a complex relationship with insecticide resistance, potentially exploiting hosts with resistance genes by increasing bacterial density, while also differentially affecting susceptibility to various insecticides, as seen with increased deltamethrin sensitivity but neutral effects on DDT resistance (32,33).

In insects, the extracellular and intracellular microbiota contribute to essential microelements beyond the host genomés metabolic capacities (34). Mosquitoes, with their life cycle comprising immature aquatic instars and one aerial adult stage, can potentially host a wide spectrum of bacteria. The microbiota plays various roles including larval nutritional support and direct impacts on adult fitness, measured as lifespan and reproduction (35–41). Despite the significant impact of microbiota on mosquitoes, there is a lack of studies on the role of maternally inherited *Wolbachia* in the fitness of *Culex* mosquitoes in the absence of their microbes. It is unclear if *Wolbachiá*s effects might be partially buffered by the microbiota. Our laboratory recently developed a method to produce germ-free *Aedes aegypti* (39). We adapted this approach to *Cx quinquefasciatus* to analyze *Wolbachia*’s role in larval development without other microbiota members.

## Results

### Development of an egg sterilization protocol for Culex quinquefasciatus

Mosquito production in sterile conditions requires protocol optimization, taking into account the specificities of mosquito species. While gnotobiotic *Ae. aegypti* (i.e. carrying a defined microbiota composition) have now been reared in several laboratories, we needed to define conditions to rear *Culex* mosquitoes in microbiologically-controlled conditions. To this aim, we used as a starting point a transient colonization method that was previously developed for *Aedes aegypti* in our laboratory, allowing to support larval development via monocolonization with a bacterium that is lost at the time of metamorphosis (39). This consists of a three-step procedure. First, eggs are surface-sterilized, producing germ-free larvae. Second, newly hatched germ-free larvae are reared in the presence of a mutant bacterium, *E. coli* HA416, which is auxotrophic for two amino-acids essential for peptidoglycan biosynthesis. As long as larvae are provided with a special sterile diet supplemented with these amino acids, *E. coli* HA416 (AUX) grows and supports larval development. Third, pupae are transferred to a rearing environment devoid of the specific amino-acids, allowing the emergence of germ-free adults. We decided to adapt this approach for field-collected *Cx quinquefasciatus* egg rafts to produce mosquitoes with no culturable microbiota.

*Ae. aegypti* eggs are laid individually and survive for weeks after drying; their hatching is stimulated when they are covered with water, notably due to the lack of oxygen. In contrast, *Cx quinquefasciatus* eggs are laid as rafts containing up to several dozens of eggs (Figure 1A), which float on the water surface. These eggs remain wet and close to the water surface throughout their uninterrupted development, limiting laboratory flexibility. When using the *Ae. aegypti* egg-sterilization protocol to surface-sterilize *Cx quinquefasciatus* eggs (Figure 1A), we observed that the egg rafts disaggregated into individual eggs or small clusters, sinking to the bottom of the flask. Almost no larvae hatched in these conditions. We hypothesized that either the eggs needed to remain attached throughout development or the raft’s location at the surface of the water was crucial for successful development, possibly due to oxygenation. To investigate the importance of the egg raft, we performed a delicate wash, minimizing raft disintegration while constantly submerging it in sterilizing solutions (sodium hypochlorite and ethanol). Alternatively, egg rafts were thoroughly washed to separate eggs, while a control group remained non-sterile. To maintain egg proximity to the surface, we incubated eggs in a minimal water layer, 1-2 mm deep (3-5 mL in a 25 cm^2^ flask laid horizontally). The next day, only 6.6% of preserved sterilized rafts hatched, compared to 71 % of individualized sterile eggs (Figure 1B-E). Non-sterile eggs exhibited the highest hatching rate (91%).

**Figure 1.**
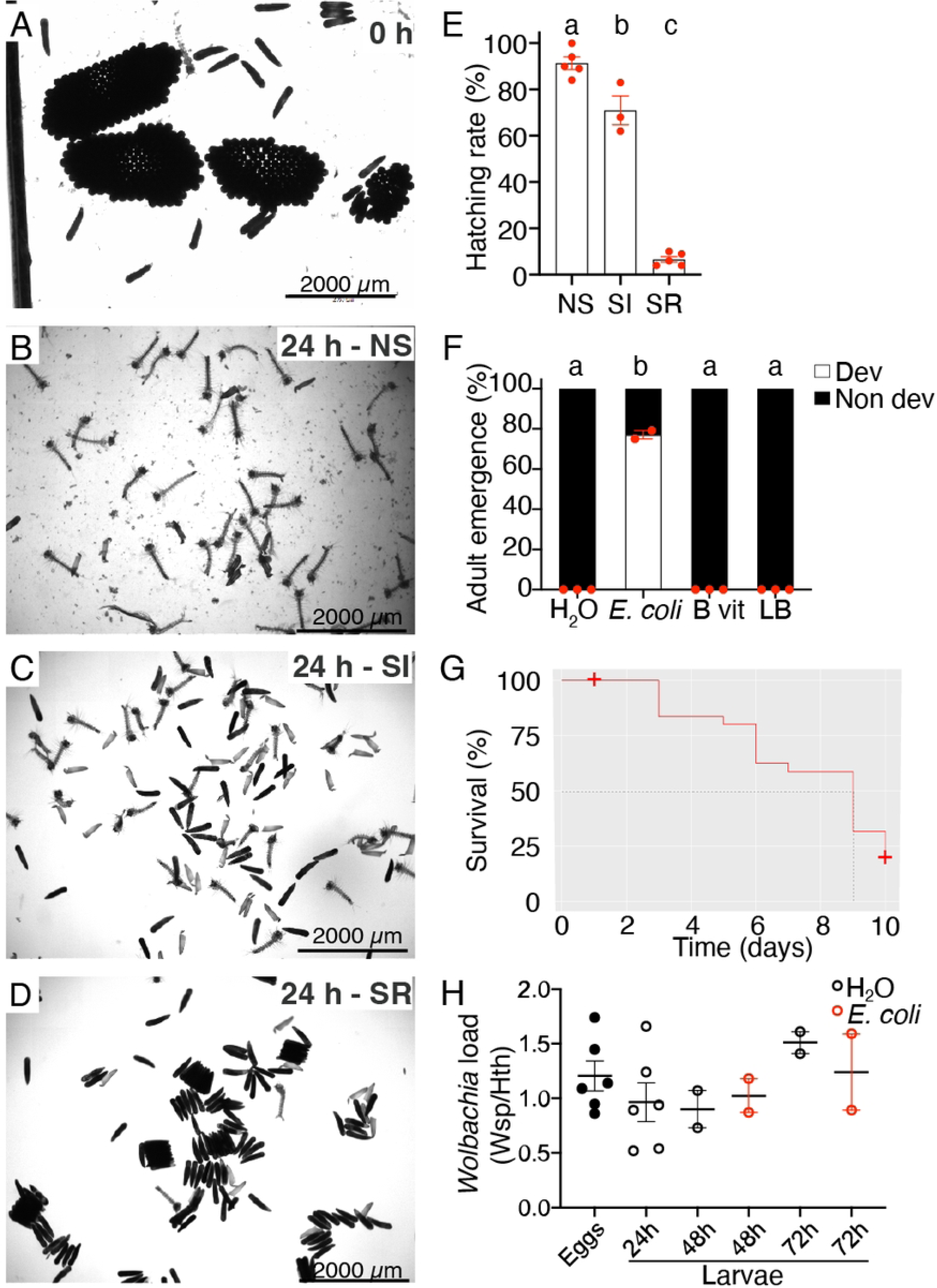
*Culex quinquefasciatus* germ-free^Wol+^ larvae hatch but do not develop without extracellular bacteria. **A.** Egg rafts collected from a local breeding site at time=0 h, just after collection. **B-D.** Aspect of the eggs 24 h after collection and treatment, whether they were not sterilized (B), sterilized with a harsh washing procedure to disrupt the rafts into individualized eggs (C) or following a delicate washing procedure, so as to preserve rafts (D). **E.** Hatching rate of the eggs imaged in B-D. NS vs SI p<0.0001, SI vs SR p<0.0001, NS vs SR p<0.0001. GLMM fit by maximum likelihood, binomial distribution, pairwise comparisons with Bonferroni adjustment. **F.** Development success of larvae on sterile diets compared to larvae monocolonized with *E. coli*. B_vit vs LB p=1.0, B vit vs Water p=1.0, B vit vs *E. coli* WT p<0.0001, LB vs Water p=1.0, LB vs *E. coli* WT p<0.0001, Water vs *E. coli* WT p<0.0001. Type III ANOVA with Satterthwaite’s method and pairwise comparisons with Bonferroni adjustment. Dev: developed to adults by day 14; Non dev: stalled in development or dead. **G.** Survival curve of sterile larvae provided with autoclaved Tetramin baby fish food. **H.** qPCR quantification of the load of *Wolbachia* in germ-free*^Wol+^* and *E. coli* colonized larvae. No difference in water vs *E. coli* p=0.77. Type III ANOVA with Satterthwaite’s method and pairwise comparisons with Tukey’s adjustment. NS – Non-sterile, SI – Sterile individuals, SR – Sterile rafts, B vit – B vitamin cocktail defined in (42), LB – lysogeny broth. In 1A-1D, representative images from 6 independent replicates. 1E, 1F, 1H; average ± SEM from 6, 3 and 2 independent replicates, respectively; 1G: average of 4 independent replicates.

The presence of *Wolbachia* in sterile larvae was confirmed by qPCR, and egg sterility was verified after incubation in liquid Luria Bertani medium (LB) or on LB agar plates. We thus refer to these individuals as germ-free*^Wol+^*, indicating they are germ-free except for the presence of the *Wolbachia* endosymbiont. To assess whether *Wolbachia* alone could support larval development, we monitored the development and lifespan of larvae provided sterile conventional food for 10 days. Larvae remained in their first instar, and supplementing the diet with a cocktail of B vitamins that are essential for *Cx quinquefasciatus* larval development (42) or with medium for bacterial culture did not improve development (Figure 1F). However, these larvae were still able to develop, as provision of live bacteria efficiently rescued 77% of individuals to adulthood. Approximately 80% of sterile larvae survived until day 5, but survival sharply declined in the following days, resulting in around 20% of surviving individuals on day 10 (Figure 1G). We confirmed by qPCR that *Wolbachia* colonization was unaffected by the absence of other bacteria. *Wolbachia* levels remained stable in first instar larvae even after 72 hours, independently of the presence of *E. coli* (Figure 1H).

Together, these data indicate that *Wolbachia* alone is insufficient to support larval development in the absence of other bacteria. Furthermore, although *Cx quinquefasciatus* typically thrives in a microbe-rich environment compared to other mosquito larvae, the sole presence of *E. coli* is sufficient to complement its requirements for larval development.

### Production of germ-free^Wol+^ Culex quinquefasciatus adults

Having successfully produced germ-free*^Wol+^* larvae, we investigated whether the transient colonization approach could yield germ-free*^Wol+^* adults. To accomplish this, we added the auxotrophic bacterial culture, using an *E. coli* wild-type as a positive control (Figure 2A).

**Figure 2.**
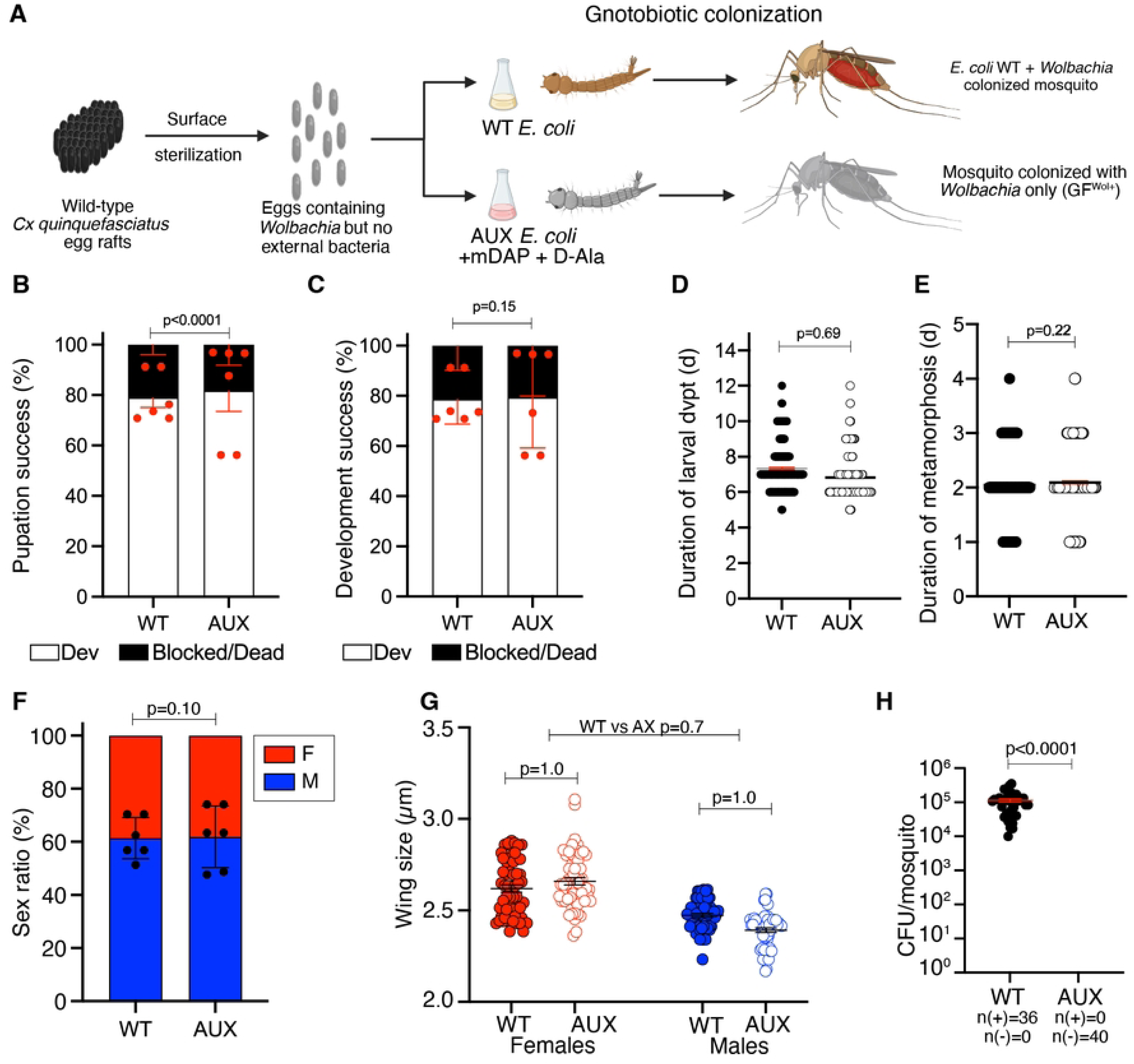
Transient colonization allows to produce germ-free*^Wol+^* adults. **A**. Experimental design for rearing germ-free*^Wol+^ or* monocolonized*^Wol+^* larvae via transient colonization. **B**. Percentage of pupation success between groups with wild-type and auxotrophic bacteria. AUX vs WT p<0.0001. GLMM fit by maximum likelihood with binomial distribution and pairwise comparison. **C**. Percentage of developmental success. AUX vs WT p=0.1432. GLMM fit by maximum likelihood with binomial distribution and pairwise comparison. **D-E.** Duration of larval development to pupae (D, AUX vs WT p<0.0001) and metamorphosis (E, AUX vs WT p=0.1767) in days. Type III ANOVA with Satterthwaite’s method and pairwise comparisons. **F**. Sex ratio in emerged adults’ post-treatment, AUX vs WT p=0.1591. GLMM fit by maximum likelihood with binomial distribution and pairwise comparison. **G.** Wing size. AUX vs WT p=1.0 Type III ANOVA with Satterthwaite’s method and pairwise comparison. **H**. Colony forming units per mosquito. AUX vs WT p<0.0001. Type III ANOVA with Satterthwaite’s method and pairwise comparison. WT - Reared with wild-type *E. coli.* AUX – Reared with auxotrophic *E. coli.* In 2B-F: average ± SEM of 6 independent replicates. 2G-H: average ± SEM of 3 independent replicates.

We monitored development daily and found no significant differences in development success rates or timing between larvae monocolonized with *E. coli* wild-type and those on AUX. Larval development success to pupa was slightly higher on AUX (Figure 2B-E). Their sex ratio and adult wing size, used as a readout of the adult size, were also similar (Figure 2F-G). Reanalyzing developmental rates data with those from subsequent experiments, we found that the overall development rate to adulthood in AUX condition increased and the development timing to pupa significantly decreased (pupae: p<0.0001, adults: p=0.18, GLMM with lsmeans), although masked here by replicate variation. Pupae transfer occurred under sterile conditions, and colony forming unit (CFU) tests confirmed that 10-day old gnotobiotic mosquitoes were colonized with *E. coli* while germ-free*^Wol+^* mosquitoes were not (Figure 2H).

### Impact of Wolbachia on development success of germ-free Culex quinquefasciatus mosquitoes

*Wolbachia* alone is insufficient to support larval development. However, previously published *Culex*-derived *Wolbachia* genomes indicate the bacterium can synthesize several B vitamins, including riboflavin, thiamin and pyridoxin, but not folate or biotin (7). Additionally, *Wolbachia* possesses complete purine and pyrimidine biosynthesis pathways. Given that bacteria-derived riboflavin and folate are essential larval development metabolites, and *Wolbachia* provides purine and pyrimidine to fly larvae (39,41,43,44), we wondered if *Wolbachia* could aid larval development under suboptimal *E. coli* conditions. Alternatively, we hypothesized that *Wolbachia* might burden larval development by consuming some nutrients. Thus, we tested the impact of *Wolbachia* load reduction on monocolonized larvae development success.

The most widely used method to clear *Wolbachia* in various insects is tetracycline antibiotic, which is also the first-choice treatment for *Culex* mosquitoes (33,45). Our AUX bacterial strain, used for transient colonization, also possesses a tetracycline-resistance cassette in addition to the kanamycin resistance cassette employed in our standardized protocol (46). We thus decided to test the impact of a tetracycline treatment on larvae colonized with AUX to assess larval development success (47). We treated larvae with three tetracycline doses (5µg/mL, 25 µg/mL, or 50µg/mL, see Methods) and confirmed that all concentrations effectively reduce *Wolbachia* load in larvae and adults (Figure 3A-B). Development outcomes were highly concentration-dependent. Treating with 5µg/mL tetracycline significantly increased development success to adulthood, indicating that *Wolbachia* is a burden rather than a nutritional benefit during larval development (Figure 3F-3G). This observation is consistent with the fact that WT-colonized larvae had a lower *Wolbachia* load and a higher development success than their AUX-colonized counterparts (Figure 3A, 3F, 3G). Increasing tetracycline concentrations progressively decreased development success, suggesting direct antibiotic toxicity to larvae (Figure 3F, 3G). The tetracycline solution at the highest concentration (50 μg/mL) underwent a noticeable color change from light yellow to brownish after 5-7 days, indicating antibiotic degradation (48). This transformation can potentially harm larvae, as tetracycline and its degradation products have been shown to have toxic effects on various organisms, including algae, at concentrations above 5 mg/mL (49).

**Figure 3.**
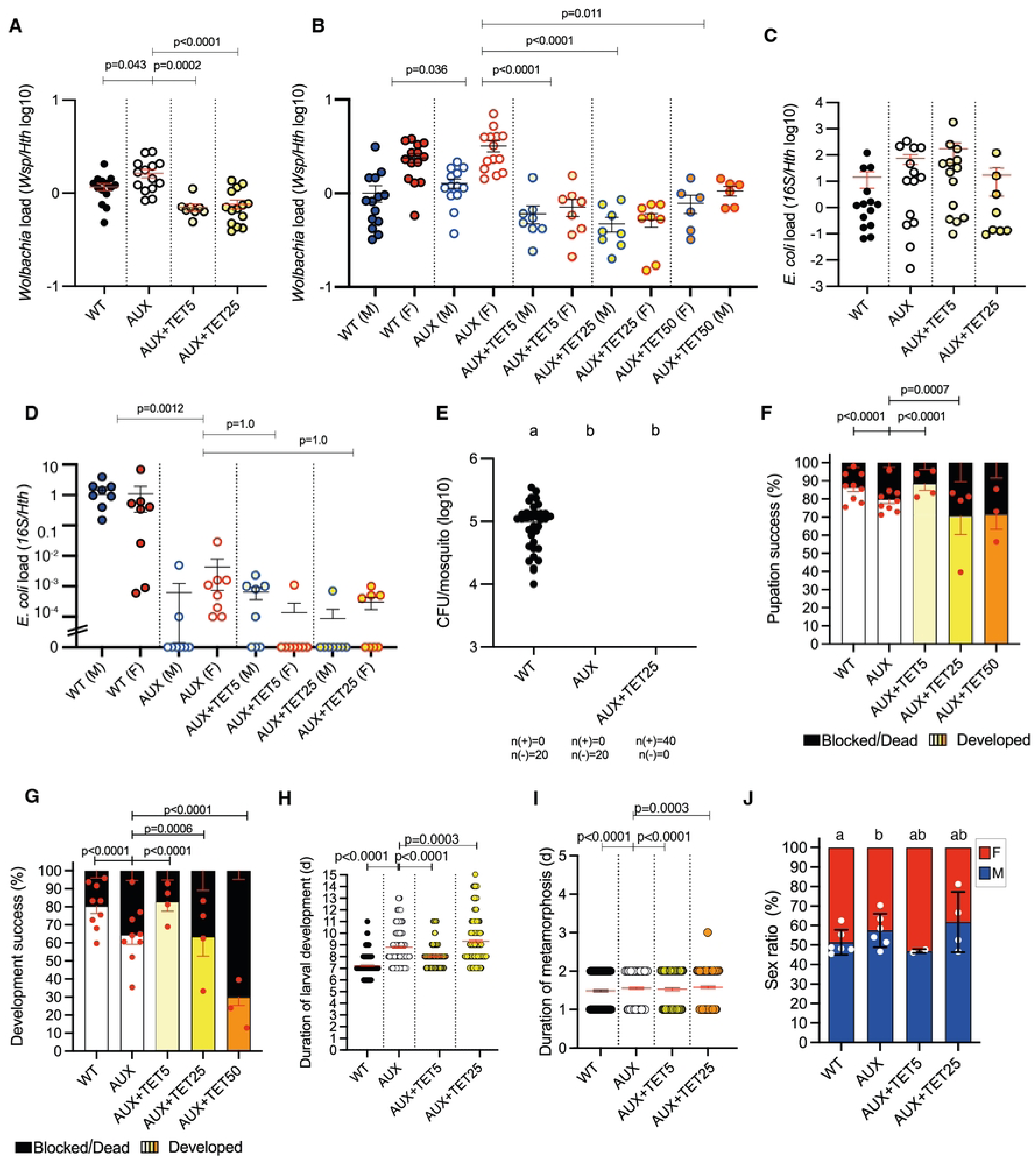
*Wolbachia* negatively impacts development success. **A.** *Wolbachia* load in larvae pools during larval development. AUX vs AUX+TET25 p<0.0001, AUX vs AUX+TET5 p=0.0002, AUX vs WT p=0.0433, AUX+TET25 vs AUX+TET5 p=1.0, AUX+TET25 vs WT p=0.0944, AUX+TET5 vs WT p=0.1790. LMM fit by REML and pairwise comparisons with Bonferroni adjustment. **B.** *Wolbachia* load in adult mosquitoes. AUX vs AUX+TET25 p<0.0001, AUX vs AUX+TET5 p<0.0001, AUX vs AUX+TET50 p=0.0111, AUX vs WT p=0.03653, AUX+TET25 vs AUX+TET5 p=1.0, AUX+TET25 vs AUX+TET50 p=1.0, AUX+TET25 vs WT p=0.0047, AUX+TET5 vs WT p=0.0221, AUX+TET50 vs WT p=0.6282. LMM fit by REML and pairwise comparisons with Bonferroni adjustment. **C.** Bacterial decolonization in larvae. AUX vs AUX+TET25 p=1.0, AUX vs AUX+TET5 p=1.0, AUX vs WT p=1.0, AUX+TET25 vs AUX+TET5 p=1.0, AUX+TET25 vs WT p=1.0, AUX+TET5 vs WT p=1.0. LMM fit by REML and pairwise comparisons with Bonferroni adjustment. **D.** Bacterial decolonization in mosquitoes. AUX vs AUX+TET25 p=1.0, AUX vs AUX+TET5 p=1.0, AUX vs WT p=0.0012, AUX+TET25 vs AUX+TET5 p=1.0, AUX+TET25 vs WT p=0.0012, AUX+TET5 vs WT p=0.0012. LMM fit by REML and pairwise comparisons with Bonferroni adjustment. **E.** Bacterial decolonization measured with CFU test. AUX vs AUX+TET5 p=1.0, AUX vs WT p<0.0001, AUX+TET5 vs WT p<0.0001. Type III ANOVA with Satterthwaite’s method and pairwise comparison with Bonferroni adjustment. **F.** Pupation ratio success in treated larvae with different tetracycline doses. AUX vs AUX+TET25 p=0.0007, AUX vs AUX+TET5 p<0.0001, AUX vs AUX+TET50 p=0.5103, AUX vs WT p<0.0001, AUX+TET25 vs AUX+TET5 p<0.0001, AUX+TET25 vs AUX+TET50 p=1.0, AUX+TET25 vs WT p<0.0001, AUX+TET5 vs AUX+TET50 p<0.0001, AUX+TET5 vs WT p=1.0, AUX+TET50 vs WT p<0.0001. GLMM fit by maximum likelihood and pairwise comparison with Bonferroni adjustment. **G.** Adult metamorphosis success in treated groups with different tetracycline doses. AUX vs AUX+TET25 p=0.0006, AUX vs AUX+TET5 p<0.0001, AUX vs AUX+TET50 p<0.0001, AUX vs WT p<0.0001, AUX+TET25 vs AUX+TET5 p<0.0001, AUX+TET25 vs AUX+TET50 p=1.0, AUX+TET25 vs WT p<0.0001, AUX+TET5 vs AUX+TET50 p<0.0001, AUX+TET5 vs WT p=1.0, AUX+TET50 vs WT p<0.0001. GLMM fit by maximum likelihood and pairwise comparison with Bonferroni adjustment. **H.** Duration of larval development until pupation. AUX vs AUX+TET25 p=0.0003, AUX vs AUX+TET5 p<0.0001, AUX vs WT p<0.0001, AUX+TET25 vs AUX+TET5 p<0.0001, AUX+TET25 vs WT p<0.0001, AUX+TET5 vs WT p<0.0001. Type III ANOVA with Satterthwaite’s method and pairwise comparison with Bonferroni adjustment. **I.** Duration of metamorphosis from pupae to adults. AUX vs AUX+TET25 p=0.0003, AUX vs AUX+TET5 p<0.0001, AUX vs WT p<0.0001, AUX+TET25 vs AUX+TET5 p<0.0001, AUX+TET25 vs WT p<0.0001, AUX+TET5 vs WT p<0.0001. Type III ANOVA with Satterthwaite’s method and pairwise comparison with Bonferroni adjustment. **J.** Sex ratio in treated groups. AUX vs AUX+TET25 p=1.0, AUX vs AUX+TET5 p=1.0, AUX vs WT p=0.0278, AUX+TET25 vs AUX+TET5 p=0.6009, AUX+TET25 vs WT p=0.0630, AUX+TET5 vs WT p=1.0. GLMM fit by maximum likelihood and pairwise comparison with Bonferroni adjustment. Average ± SEM of 3 (3A, 3E), 4 (3B, 3C) and 2 (3D), independent replicates. In 3F-3J, average ± SEM of 9 independent replicates (12 replicates), treated groups independent replicates were 4 in AUX+TET5 and AUX+TET25 conditions, and 3 for AUX+TET50 (F-J).

Development of AUX-colonized larvae and pupae was delayed compared to WT-colonized ones; this was partly rescued in 5 µg/mL tetracycline treated group (Figure 3H-3I). Similarly, AUX-colonized larvae led to a slightly lower proportion of females than WT-colonized ones and 5 µg/mL tetracycline-treated ones. As females require more nutrients for development than males, this points to a lower nutritional status in AUX conditions. CFU analysis confirmed that WT-colonized mosquitoes indeed contained live *E. coli,* while the AUX and AUX+TET did not (Figure 3E), which matches the loss of bacteria after metamorphosis while not during larval stages (Figure 3C-D).

## Discussion

In this study, we established a protocol to produce mosquitoes harboring solely *Wolbachia* and no other microbiota. We demonstrated that *Wolbachia* alone is insufficient to support larval development. Conversely, our data indicate that this endosymbiont negatively impacts mosquito larval development.

*Culex* mosquitoes are generally known to grow in “dirty” water, i.e. stagnant bodies of water that are turbid (1,50,51), whereas *Aedes* mosquitoes typically breed in cleaner water or with a moderate amount of organic matter present (52). For instance, we generally set *Aedes* oviposition traps with clear water, while we add chicken manure to attract *Culex.* Hence, one may expect that *Cx quinquefasciatus* larvae require more bacteria, and potentially a higher diversity of microorganisms, to grow compared to *Aedes aegypti*. However, we observed that *E. coli* alone was able to rescue larval development of *Cx quinquefasciatus,* as already found for *Ae. aegypti*. Nonetheless, we achieved a slightly lower success rate (∼75 %) than what we typically observe with *Ae. aegypti* (∼85%), particularly when rearing mosquitoes on AUX *E. coli* (∼60%), which tends to be present at lower loads at the end of larval development (39). This suggests that there might still be a difference in the requirement for microbial metabolites between the two mosquito species, which may be quantitative rather than qualitative. Moreover, we observed that *E. coli* is highly efficiently transmitted through metamorphosis in *Cx quinquefasciatus*, as 100% adults reared on WT-*E coli* were positive to the bacterium, compared to only 50% in *Ae. aegypti* (Figure 2H and (39)).

The eggs of *Culex* mosquitoes are laid as a raft, allowing them to remain on the water surface. Consequently, their upper side is dry and in direct contact with open air. Our rigorous sterilization protocol caused the eggs to sink into the hatching water, and a low hatching rate was observed only when the water layer was very thin and the eggs were individualized. We interpret these results as indicating a high oxygen requirement for *Culex* egg development (53–55).

The presence of *Wolbachia* in a high proportion of terrestrial and aquatic insects is due to its ability to manipulate reproduction through cytoplasmic incompatibility, its stability within the insect, and its efficient vertical transmission. Our data corroborate this stability in *Cx quinquefasciatus*, as the antibiotic treatment did not allow complete clearance of *Wolbachia.* In line with these observations, protocols for antibiotic treatments to clear *Wolbachia* generally involve treatment on several generations (45). Beyond its efficient colonization through manipulation of mosquito reproduction, *Wolbachia*’s ability to colonize populations can be enhanced by positively impacting mosquito fitness. Symbionts combining parasitism and mutualism to enhance their population colonization are often termed “Jekyll and Hyde” symbionts (56), in reference to the *Dr Jekyll and Mr Hyde* novel, whose main character recurrently shifts between being a criminal and a gentleman. *Wolbachia* has notably been shown to provide biotin and riboflavin to bedbugs (8,12) and planthoppers (13). It has recently been reported as a nutritional symbiont in *Drosophila*, shifting to an energy-save profile during growth and development, and increasing resistance to stress (43).

As microbiota-derived B vitamins, especially riboflavin, are crucial for mosquito development (41), we hypothesized that *Wolbachia* might also support larval development in *Culex sp.*, which are heavily colonized by *Wolbachia*. The genome of *Culex*-associated *Wolbachia*, wPip, lacks the biotin operon but is predicted to possess a complete riboflavin biosynthesis pathway (13). However, our data indicate that *Wolbachia* clearance benefits mosquito development, suggesting that *Wolbachia* is detrimental to larval development under our experimental conditions. We hypothesize that *Wolbachia* competes with its host for resources. Our findings align with a previous study showing a non-significant trend towards lower development success of *Wolbachia*-colonized compared to *Wolbachia*-cured *Aedes albopictus* larvae at a medium larval density of 1 larva/mL (57).

These data indicate that *Wolbachia* is a metabolic parasite during larval development, in addition to its previously characterized role as a reproductive parasite. Nevertheless, it may still be considered a Jekyll and Hyde symbiont, as it can benefit its host inducing resistance to viruses in flies and mosquitoes (58). This protective activity is suggested to enhance *Wolbachia*’s success. Indeed, *Wolbachia*-mediated antiviral protection has been shown to prolong survival in various host-virus combinations, thereby positively impacting host fitness (59). Furthermore, *Wolbachia* has been found to protect *Drosophila* against viral infections at 25°C but not at 18°C, correlating with higher *Wolbachia* colonization in tropical compared to temperate regions (59).

Our data further indicates a concentration-dependent toxicity of tetracycline in mosquito larvae. Tetracycline inhibits bacterial protein synthesis via interacting with the 30S ribosomal subunit, inhibiting the binding of aminoacyl-tRNAs (60). While ribosomes of eukaryotes, including in insects, are composed of 60S and 40S subunits and are hence not sensitive to tetracyclines, mitochondrial ribosomes may be targeted by this drug (61). So far, several negative impacts of tetracycline on insects have been reported, yet some may be linked to the loss of the microbiota. A tetracycline treatment to clear *Culex pipiens* of *Wolbachia* using 10-20% solutions resulted in a development success as low as 10-15% (62), yet this strong impact may rather be due to the loss of other microbes, which are essential for mosquito larval development. Some other studies on *Wolbachia* clearance in *Culex* larvae did not report survival rates or indicated no effect on larval development, which may be linked to the presence of tetracycline-resistant bacteria (30,45,47,63). In ticks, tetracycline has been found to reduce the reproductive fitness by downregulating vitellogenesis and embryogenesis, although this effect is likely linked to microbiota depletion (64). Our results obtained in otherwise germ-free conditions, indicate a direct toxic impact of tetracycline on mosquitoes. They are consistent with a previous report of long-lasting impact of tetracycline on mitochondrial metabolism (65).

Beyond arthropods, tetracycline has been documented to cause DNA damage, metabolic alterations, and oxidative stress in earthworms (66). In zebrafish embryos, exposure to this antibiotic also leads to developmental delay, reactive oxygen species (ROS) production, and cell death (67). In common freshwater green algae, the active form (tetracycline) and most common degraded forms (anhydrotetracycline and epitetracycline) inhibited the growth of *Chlamydomonas reinhardtii*, with tetracycline exhibiting the highest toxicity (49). We observed a brown staining after a few days in our larval breeding water, indicating that it becomes oxidized over time. It remains unclear whether the toxicity observed in our larvae can be attributed to tetracycline or to its oxidized product.

## Conclusions

While *Wolbachia* studies related to the impact in fitness in mosquitoes are sometimes hindered by the fact that antibiotic treatments affect the insect microbiota, we managed to set up a method to investigate the role of *Wolbachia* in *Culex* mosquitoes in the absence of other microbiota members. We observed that *Wolbachia* is not able to support larval development on its own. This is consistent with the absence of the biotin operon reported in *Wolbachia pipiens*, contrary to strains found in bedbugs and planthoppers. Our results indicate that *Wolbachia* has a parasitic impact on larval development, hence point that reproductive parasitism in itself is sufficient to support the strong colonization capacity of *Wolbachia* in *Cx quinquefasciatus* populations.

## Materials and Methods

### Culex quinquefasciatus mosquitoes

Up to three breeding sites were placed inside the campus of the Institut Pasteur de la Guyane in Cayenne, French Guiana (GPS coordinates: 4.943100, -52.325350) for collecting egg rafts. They consisted in containers containing tap water, soil and chicken manure, as this organic matter-enriched water attracts wild gravid *Cx quinquefasciatus* females to lay eggs. Each breeding site was checked daily, covered with a net on weekends and changed every 14 days to avoid potential escaping mosquitoes from uncollected larvae. Mosquito species were confirmed using a taxonomy key at the beginning of the project (68,69). As this species is by far the commonest species in urban areas in French Guiana (69), further checks were not performed in subsequent experiments, but no variations in the shape of egg rafts, larvae or adults were noticed during stereoscopic observations.

### Sterilization protocol and bacterial inoculum

Egg rafts were recovered from the breeding site bucket and transported to the lab, a pre-wash with tap water was made to remove organic matter from the liquid. Under the biosafety cabinet rafts were sterilized following lab protocol (39). Sterile eggs were kept in sterile conditions and placed in a 25 cm^2^ culture flask with a minimal water layer (∼3-5 mL depending on the number of eggs) in horizontal position, a sterility control test in a petri dish with LB agar was tested each experiment and kept until 48 h to verify sterility.

After egg rafts sterilization protocol, 24 h post-hatched first instar larvae were individually placed into 24-wells plates with 1.5 mL of *E. coli* HA416 supplemented with D-Alanine (50 µg/mL) and acid meso-diaminopimelic (12.5 µg/mL), with one drop (0.05-0.1 mL) of sterile 5 % (w/v) TetraMin baby™ fish food solution. Plates were maintained in a climatic chamber at 28 °C with a light-dark cycle 12 h/12 h with 80 % relative humidity.

For a control rearing, *E. coli* HS (wild-type, WT) was used in the same culture conditions as the mutant bacteria. Sterile larvae survival, pupation and development success tests, were recorded daily during 14 days. Hatching rates were counted under microscope at 24 h after sterilization.

### Antibiotic treatment during transient colonization rearing

*E. coli* HA416 contains a tetracycline resistance cassette, a parallel treatment with the antibiotic to remove *Wolbachia* was viable without hindering bacterial culture added to rear axenic mosquitoes. An initial dose of 5, 25 and 50 μg/mL was added to each larva, and then at day 3 and 5, 25 and 50 μg/mL were added again to sustain a level on the antibiotic concentration, also at day 2 and 4 bacterial culture was added as well in “AUX” treatment (33,45).

### Wolbachia detection

For detection of *Wolbachia,* individual female and male mosquitoes were placed in screwcaps tubes with crystal beads and frozen at -80 °C until use. Samples were processed for DNA extraction using the MagBio HighPrep Blood and Tissue DNA kit. Tissue lysis was performed with a Precellys Evolution bead beater homogenizer (Berting Technologies) following manufacturer’s conditions.

Protein digestion was completed with Proteinase K incubation overnight at 55 °C. Automatic DNA extraction was accomplished with the kit in a KingFisher Duo Prime system (Thermo Scientific). DNA was eluted in nuclease-free water and saved at -80 °C until use. Genomic DNA was diluted to 25 ng/μL using a Nanodrop spectrophotometer (Thermo Scientific) for qPCR test. Quantitative PCR test was performed in a Roche 480 Lightcycler with a 2x SYBR-green master mix using standard protocol instructions (Takara). Total *Wolbachia* density was analyzed by relative quantification of the *Wolbachia surface protein* against the mosquito *homothorax* gene. *Wolbachia* primers, *wsp* (Fwd) 5’ - ATCTTTTATAGCTGGTGGTGGT - 3’, *wsp* (Rev) 5’ - AAAGTCCCTCAACATCAACCC -3’; housekeeping *hth* (Fwd) 5’-TGGTCCTATATTGGCGAGCTA – 3’, *hth* (Rev) 5’-TCGTTTTTGCAAGAAGGTCA – 3’ (Ant et al., 2020).

### Statistical analysis

Egg rafts and individual first instar larvae were photographed with a stereomicroscope.

Numerical observation data was collected and organized in excel sheets, statistical analysis was performed in R studio 2023.06.2+561 under R 4.3.1. Survival curve was made using survival and survminer packages. Multicomparisons were performed using genealized mixing models (GLMM) with the packages lsmeans (70), and glmer (71).

Bar and scatter plots were made using GraphPad Prism 10.3.1 for Mac.

## Acknowledgements

We would like to thank Jean-Géraud Issaly, deceased in 2022, who contributed to the preparation of the capture site and the identification of the mosquito species during the beginning of the project. We thank Patrick Mayen for providing the chicken manure for the preparation of our capture site.

